# DNA methylation change in neurotrophic genes with aging and delirium evidenced from three independent cohorts

**DOI:** 10.1101/730382

**Authors:** Taku Saito, Patricia R. Braun, Sophia Daniel, Sydney S. Jellison, Mandy Hellman, Eri Shinozaki, Sangil Lee, Hyunkeun R. Cho, Aihide Yoshino, Hiroyuki Toda, Gen Shinozaki

## Abstract

**INTRODUCTION:** We previously reported the association between DNA methylation (DNAm) of pro-inflammatory cytokine genes and aging. Neurotrophic factors are also known to be associated with aging and neurocognitive disorders. Thus, we hypothesized that DNAm of neurotrophic genes change with aging, especially in delirium patients.

**METHODS:** DNAm were analyzed using HumanMethylationEPIC BeadChip Kit in 3 independent cohorts; blood from 383 Grady Trauma Project subjects, brain from 21 neurosurgery patients, and blood from 87 inpatients with and without delirium.

**RESULTS:** Both blood and brain samples showed that most of the DNAm of neurotrophic genes were positively correlated with aging. Furthermore, DNAm of neurotrophic genes were positively correlated with aging in delirium cases than in non-delirium controls.

**DISCUSSION:** These findings support our hypothesis that the neurotrophic genes may be epigenetically modulated with aging, and this process may be contributing to the pathophysiology of delirium.

## INTRODUCTION

Although delirium is common among elderly patients [1], it is underdiagnosed and undertreated [2]. Various screening methods have been developed to characterize the epidemiology and risk factors of delirium (e.g., the Confusion Assessment Method (CAM) (Inouye et al., 1990) and Confusion Assessment Method for Intensive Care Unit (CAM-ICU) [3, 4]). Although they have excellent sensitivity and specificity in research settings, they are found to have suboptimal sensitivity (38–47%) in the context of “real world” intensive care units [5, 6]. Thus, among the elderly at high risk for delirium, biomarkers of delirium would aid in the prediction and treatment of delirium.

Previously we reported the association of DNA methylation (DNAm) changes in pro-inflammatory cytokine genes with aging, indicating its possible role in the pathophysiology of delirium [7]. This emphasized past findings of the importance of neuroinflammation in the development of delirium with the additional link of DNAm changes. However, the etiology of delirium is complex with many facets leading up to its manifestation, and it is important to look at additional factors involved. Given the evidence of neurotrophic factors being associated with age-related behavior, it is possible that DNAm of these genes may be differentially regulated in the context of aging with delirium [8, 9].

Neurotrophic factors are signaling molecules that contribute to neural plasticity, learning, and memory [10]. Decreased serum levels of the brain-derived neurotrophic factor (BDNF) were associated with cognitive impairment in elderly people [11]. Serum levels of the glial cell line-derived neurotrophic factor (GDNF) as well as BDNF were decreased in subjects with mild cognitive impairment and Alzheimer’s disease [12]. A Neuronal Per-Arnt-Sim domain protein 4 (NPAS4) knockdown mouse model showed increased cell death in cortical neurons [13]. This *in-vivo* investigation also revealed an increased lesion size and greater neurodegeneration after a photochemically-induced stroke [13]. The nuclear receptor subfamily 4A2 (NR4A2) critically regulates Alzheimer’s disease (AD)-related pathophysiology [14]. A *NR4A2* knockdown mouse model exacerbates AD symptoms/pathology of neuroinflammation/degeneration, and plaque accumulation. Administration of the FDA-approved *NR4A2* modulating drug significantly reduced cognitive impairments and plaque numbers [14].

Aging alters gene expression in the brain drastically [15, 16] and such changes are regulated by a variety of epigenetic modifications including DNAm [17], histone modifications [18], and micro RNA-mediated transcriptional control [19, 20]. Indeed, DNAm patterns have been shown to change dynamically throughout the human lifespan [21], and epigenetic processes involved in aging have been studied in detail [22–26]. It has previously been shown that DNAm levels of *BDNF* is significantly correlated with age in blood [27], but whether this is exacerbated or dysregulated in age-related disorders, such as delirium, needs to be studied. Our aim was to first expand upon this study and examine whether DNAm of additional neurotrophic genes are significantly correlated with aging in blood and brain tissues in non-delirium populations. Second, we wanted to compare to what degree the DNAm levels of the neurotrophic genes were correlated in delirium cases versus controls.

In this study, we analyzed samples from three cohorts to examine the degree of correlation between DNAm and age in neurotrophic genes. We hypothesized that DNAm in neurotrophic genes in both brain and blood would increase with aging, and that these associations would be more significant in delirium patients.

## METHODS

### Grady Trauma Project (GTP) Cohort Sample Processing

Three hundred and eighty-three subjects (age average = 41.50, age SD = 12.93, age range = 18-77, female; n = 273, race = 380 African American, 2 Native American and 1 other) were analyzed from the Grady Trauma Project (GTP) cohort. Detailed information of this cohort has been previously described [7]. DNAm data was processed by using Illumina HumanMethylation450. Processing of the dataset has been described previously [28].

### Neurosurgery (NSG) Cohort Sample Processing

Twenty-one subjects were analyzed from the neurosurgery (NSG) cohort. Details of this cohort are described in a previous study [29]. Resected brain tissue was collected from subjects with intractable epilepsy who underwent neurosurgery. Written informed consent was obtained from all of the participants. This study was approved by the University of Iowa’s Human Subjects Research Institutional Review Board. DNAm was assessed with the Infinium HumanMethylationEPIC BeadChip™ Kit (Illumina, WG-317-1002). The R packages RnBeads and Minfi [30, 31] were used to process the raw data and perform quality control checks, data filtering, and normalization techniques [30–32].

### Epigenetics of Delirium (EOD) Cohort Subjects

A separate ongoing study of delirium was performed at the University of Iowa Hospitals and Clinics. Further characteristics of this cohort were detailed previously [7, 33, 34]. Ninety-two subjects participated between November 2017 and October 2018. Among them, eighty-seven subjects (age average = 70.2 years, age SD = 10.2, age range = 42–101 years, male; n = 60) were analyzed as described below. Written informed consent was obtained from all of the participants, and this study was approved by the University of Iowa’s Human Subjects Research Institutional Review Board.

### Definition of Delirium Status

Details of the EOD cohort inclusion/exclusion criteria have been previously described [33]. Briefly, participants were screened for delirium by hospital records, the Confusion Assessment Method for Intensive Care Unit (CAM-ICU) [3], the Delirium Rating Scale - Revised-98 (DRS-R-98) [35], and the Delirium Observation Screening Scale (DOSS) [36]. A final decision of delirium phenotyping was conducted by trained psychiatrist (G.S.).

### EOD Sample Processing

For the EOD cohort, whole blood samples were collected with EDTA tubes, and all samples were stored at −80 °C. Genomic DNA was extracted with the MasterPure™ DNA Purification kit (Epicentre, MCD85201) and bisulfite-converted with the EZ DNA Methylation™ Kit (Zymo Research, D5002).

### EOD Methylome Ananalysis

The Infinium HumanMethylationEPIC BeadChip™ Kit (Illumina, WG-317-1002) was used to process DNAm levels of 93 samples (two samples were from one subject). The R packages ChAMP [37] and minfi [30, 38] were used to process the raw data. Using the ChAMP package, one sample was filtered out because it had above 10% of sites with detection p-value greater than 0.01, and CpG sites were filtered out as described below. This resulted in 92 samples and 701,196 probes. Additional outlier samples were found with the density and multidimensional scaling plots. This included five samples, two of which were the duplicate samples from the same individual. The remaining 87 samples (43 samples were cases) were reprocessed with the ChAMP package. CpG sites were filtered out if they 1) had a detection ρ-value greater than 0.01 (12,903 probes), 2) had less than three beads in at least 5% of samples per probe (8,280 probes), 3) were non-CpG probes (2,911 probes), 4) were SNP-related probes [39] (94,425 probes), 5) were cross-reactive probes [40] (11 probes), or 6) were located on the X or Y chromosome (16,356 probes).. Samples were normalized with beta mixture quantile dilation [41], and the Combat method was used to correct for batch effect [42, 43]. The final dataset contained 731,032 sites.

### Statistical Analysis

All statistical analyses were performed with R [44]. The correlation between age and DNAm levels of neurotrophic genes in each CpGs were tested by Pearson’s correlation analysis.

## RESULTS

### DNA methylation and expression from GTP cohort blood samples

We investigated the association between aging and DNAm levels at CpGs in seven neurotrophic genes; *BDNF, GDNF, activity regulated cytoskeleton associated protein* (*ARC*), *Fos Proto-Oncogene, AP-1 Transcription Factor Subunit (FOS), NPAS4, nuclear receptor subfamily 4A1 (NR4A1) and NR4A2*. To test this, we first used the GTP cohort that includes blood samples from 383 subjects. In the processed GTP methylome dataset, there are 226 CpGs annotated to be in the seven neurotrophic genes. Of these, DNAm of 16 CpGs were significantly associated with aging genome-wide significance levels (p < 5 × E-8), including seven CpGs in *BDNF* and four CpGs in *GDNF*. The top hits were at a CpG in in *GDNF* (cg02328239; p = 1.54 × E-20), and a CpG in *BDNF* (cg05733135; p = 4.49 × E-15) (Table 1, Figure 1). Among top 53 CpGs with p < 2 × E-4 (Bonferroni corrected for 226 CpGs, significant level p < 0.05/226), 49 CpGs (92.5%) were positively correlated with aging (Table 1). 109 CpGs were associated with aging at nominal significance (p < 0.05). Correlations between age and DNAm levels at 226 CpGs were shown in Supplementary Table 1.

**Figure 1.**
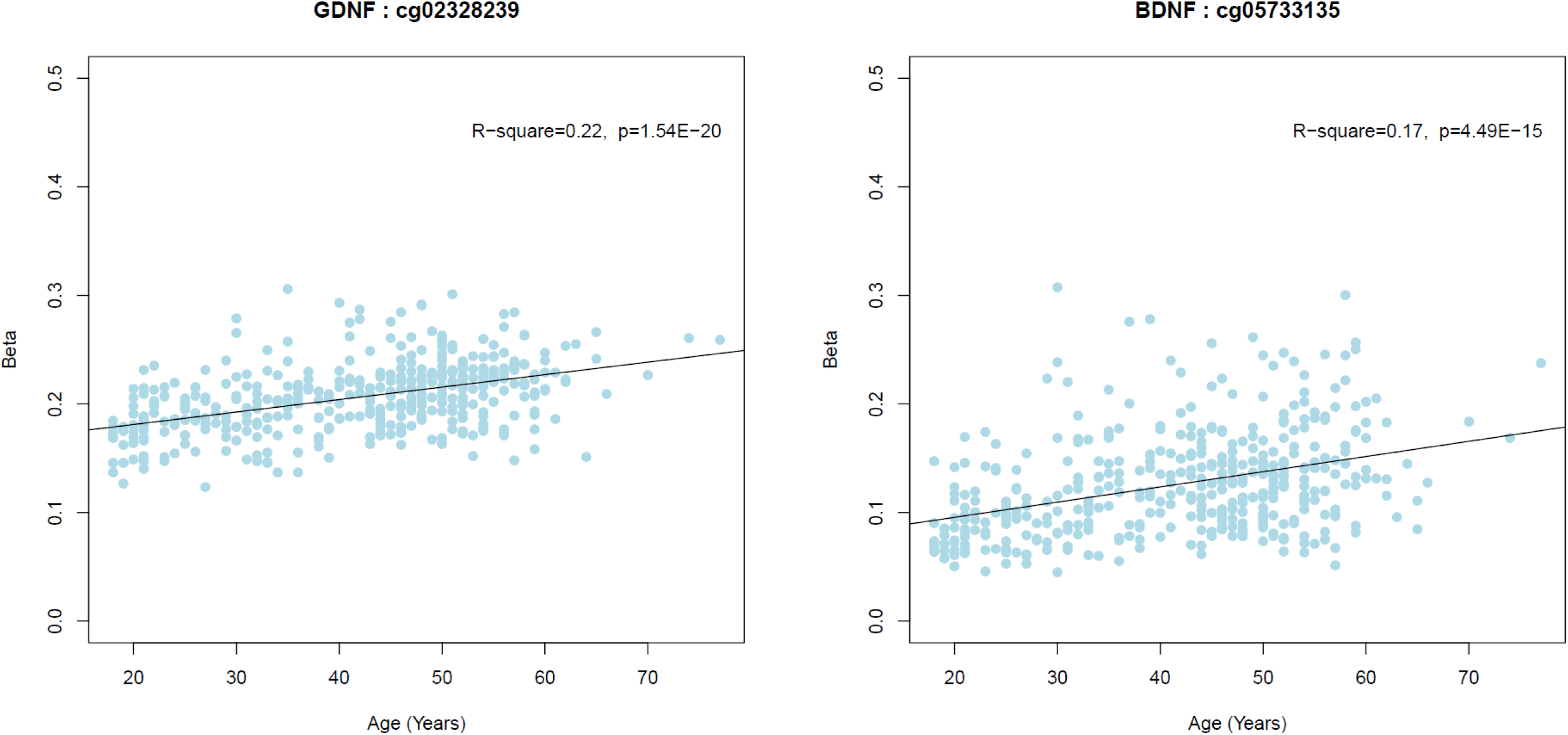
Correlation between age and beta value in the top 2 CpGs. Abbreviations: BDNF; Brain-derived neurotrophic factor, GDNF; glial cell-derived neurotrophic factor.

**Table 1:**
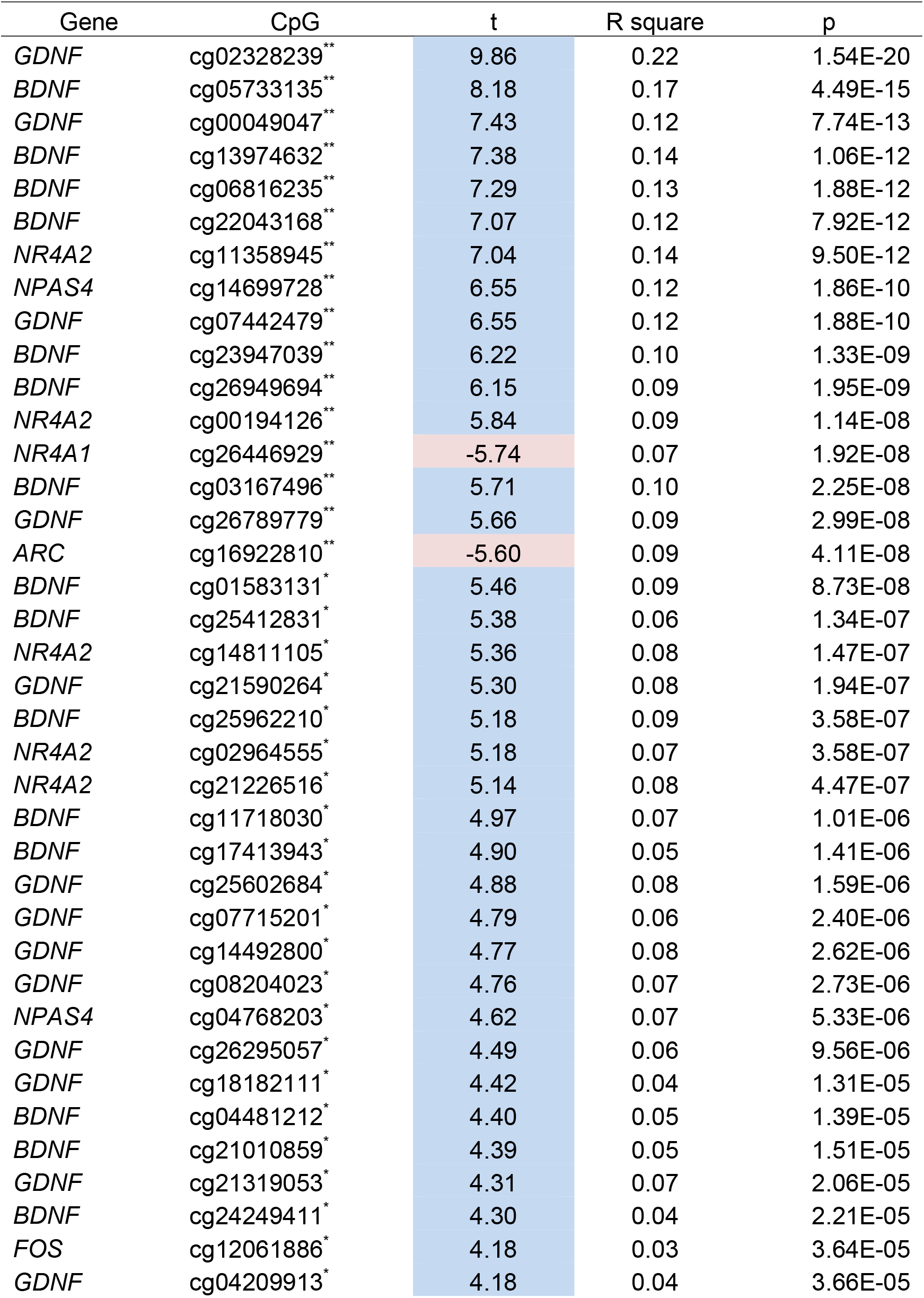
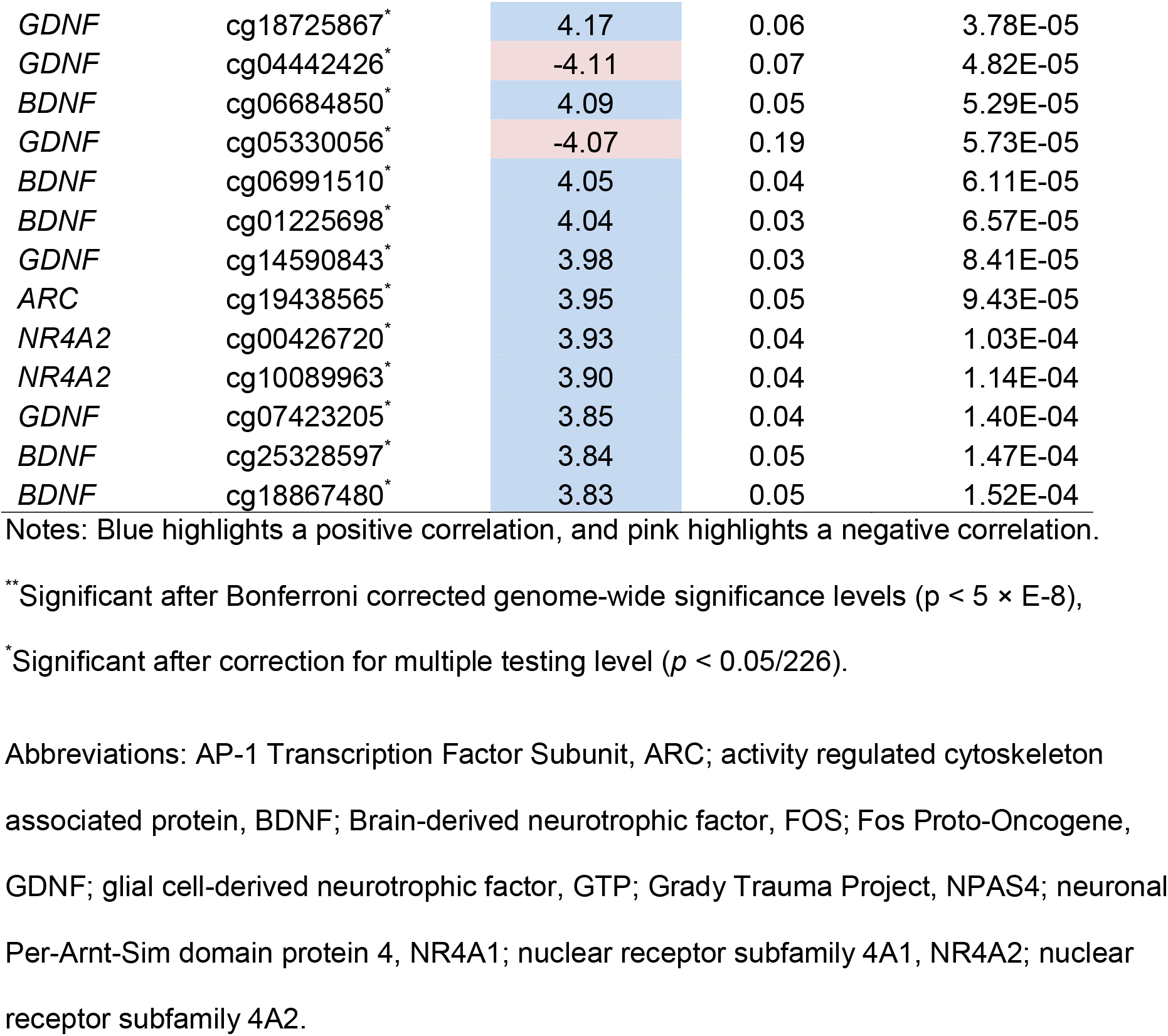
Correlations between age and DNAm levels of neurotrophic genes in blood samples obtained from the GTP cohort

### NSG study samples, DNAm in brain

We next tested whether these associations of DNAm in the GTP cohort were also present in brain tissue in our NSG cohort (N = 21). We focused on top 4 genes based on the 16 CpGs significantly associated with aging at genome-wide significant levels in the GTP cohort (*BDNF, GDNF, NR4A2*, and *NPAS4*). In *BDNF*, 18 CpGs were positively correlated with aging at nominal significance (p < 0.05), whereas 4 CpGs were negatively correlated with aging at nominal significance (Table 2). Similarly, 15 CpGs were positively correlated with aging at nominal significance, whereas no CpGs were negatively correlated with aging at nominal significance in *GDNF*, 20 CpGs were positively correlated with aging at nominal significance, whereas 3 CpGs were negatively correlated with aging at nominal significance in *NR4A2*, and 2 CpGs were positively correlated with aging at nominal significance, whereas no CpGs were negatively correlated with aging at nominal significance in *NPAS4* (Table 2). Furthermore, the top hit CpG at cg02328239 in *GDNF* in the GTP study (p = 1.54 × E-20) was also positively correlated with aging at multiple testing significant level (Bonferroni corrected for 201 CpGs, significant level p < 0.05/201 = 2.49 × E-4) in brain tissue in the NSG study (Table 2). In fact, among the nominal significant (p < 0.05) CpGs in brain tissue in the NSG study, 3 CpGs in *BDNF*, 3 CpGs in *GDNF*, 3 CpGs in *NR4A2*, and 2 CpGs in *NPAS4* were also nominally significant in the GTP study (Table 1, 2). Correlations between age and DNAm levels at 201 CpGs were shown in Supplementary Table 2.

**Table 2:**
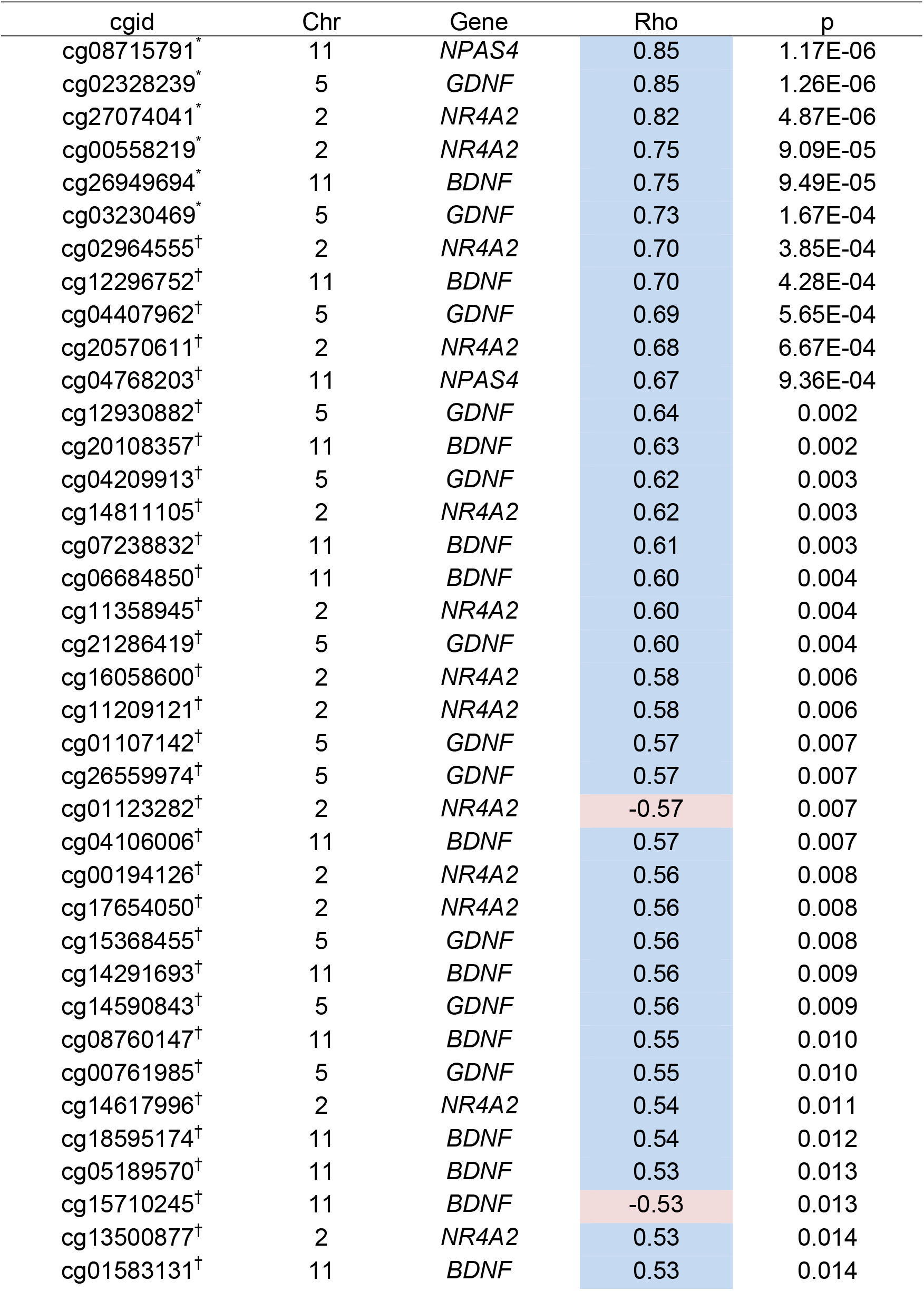
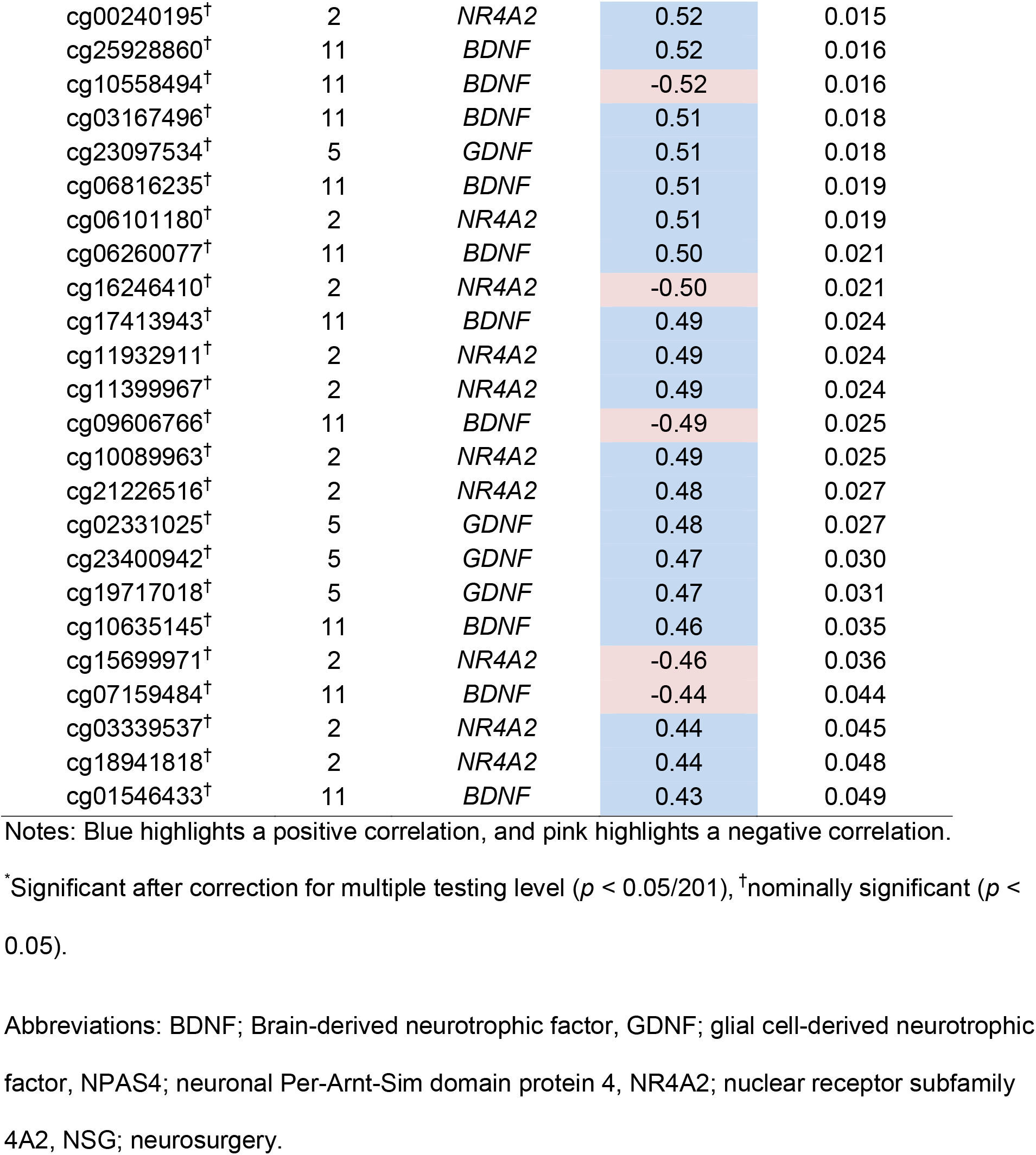
Correlations between age and DNAm levels of neurotrophic genes in brain samples obtained from the NSG cohort

### Comparison of delirium cases vs non-delirium controls in EOD study samples

We next expanded upon our previous analyses in population of delirium patients to determine if there were differences in the degrees of correlation for neurotrophic genes between DNAm and aging among delirium cases and controls. We evaluated the degree of correlation between age and DNAm levels at CpGs in *BDNF* (Table 3), *GDNF, NR4A2*, and *NPAS4* (Supplementary Tables 3-5). Delirium cases showed 1 significant CpG after correction for multiple testing level (p < 0.05/192 = 7.35 × E −4) in *BDNF*, whereas non-delirium controls showed no significant CpGs after correction for multiple testing level (Table 3). Furthermore, there were more nominal significant (p < 0.05) CpGs in delirium cases than in non-delirium controls with *BDNF* (5 in delirium cases vs. 3 in non-delirium controls), *GDNF* (5 vs. 2), and *NR4A2* (3 vs. 0). Also, there were more positively correlated CpGs in delirium cases than in non-delirium controls with *BDNF, GDNF*, and *NPAS4*.

**Table 3:**
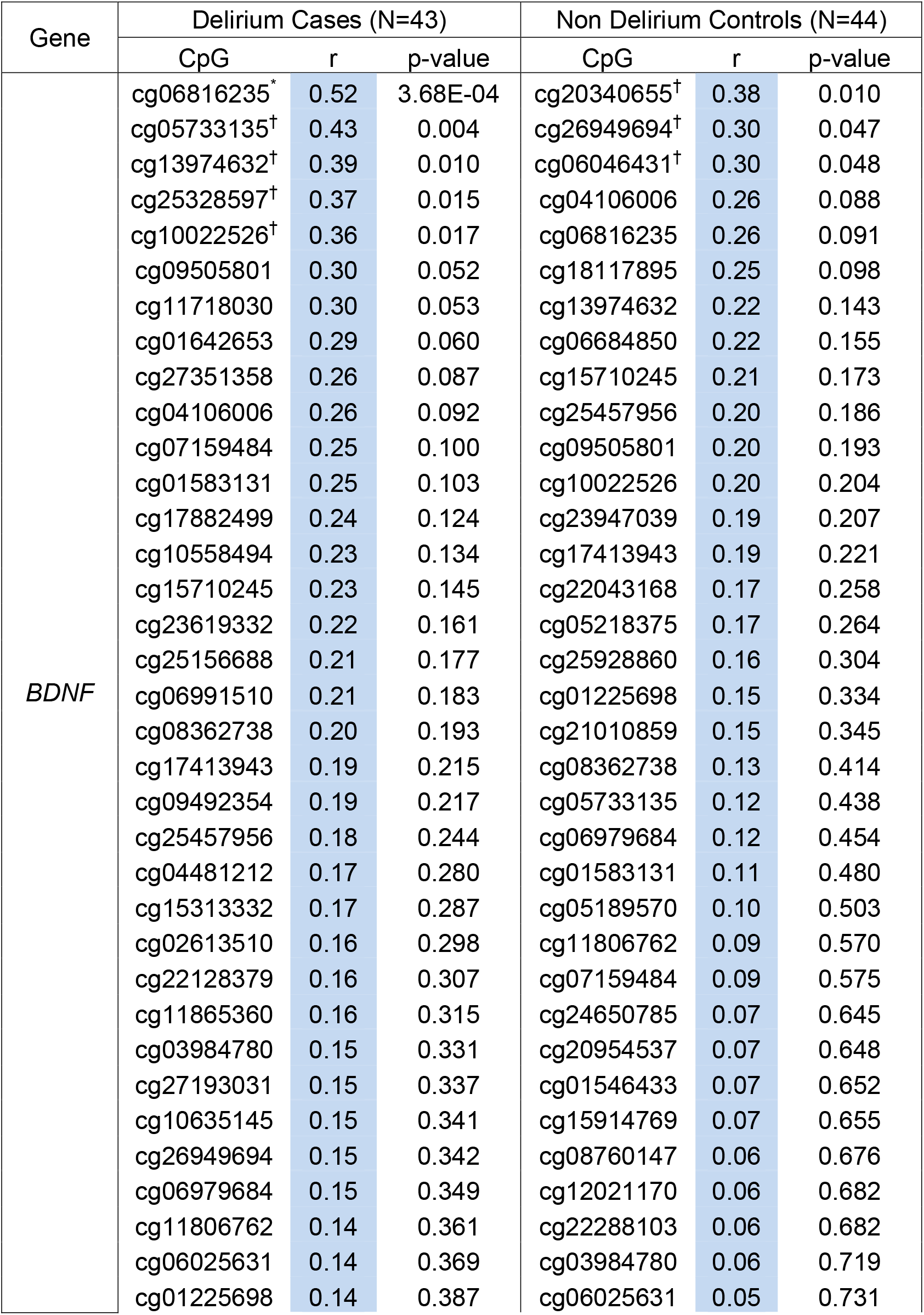
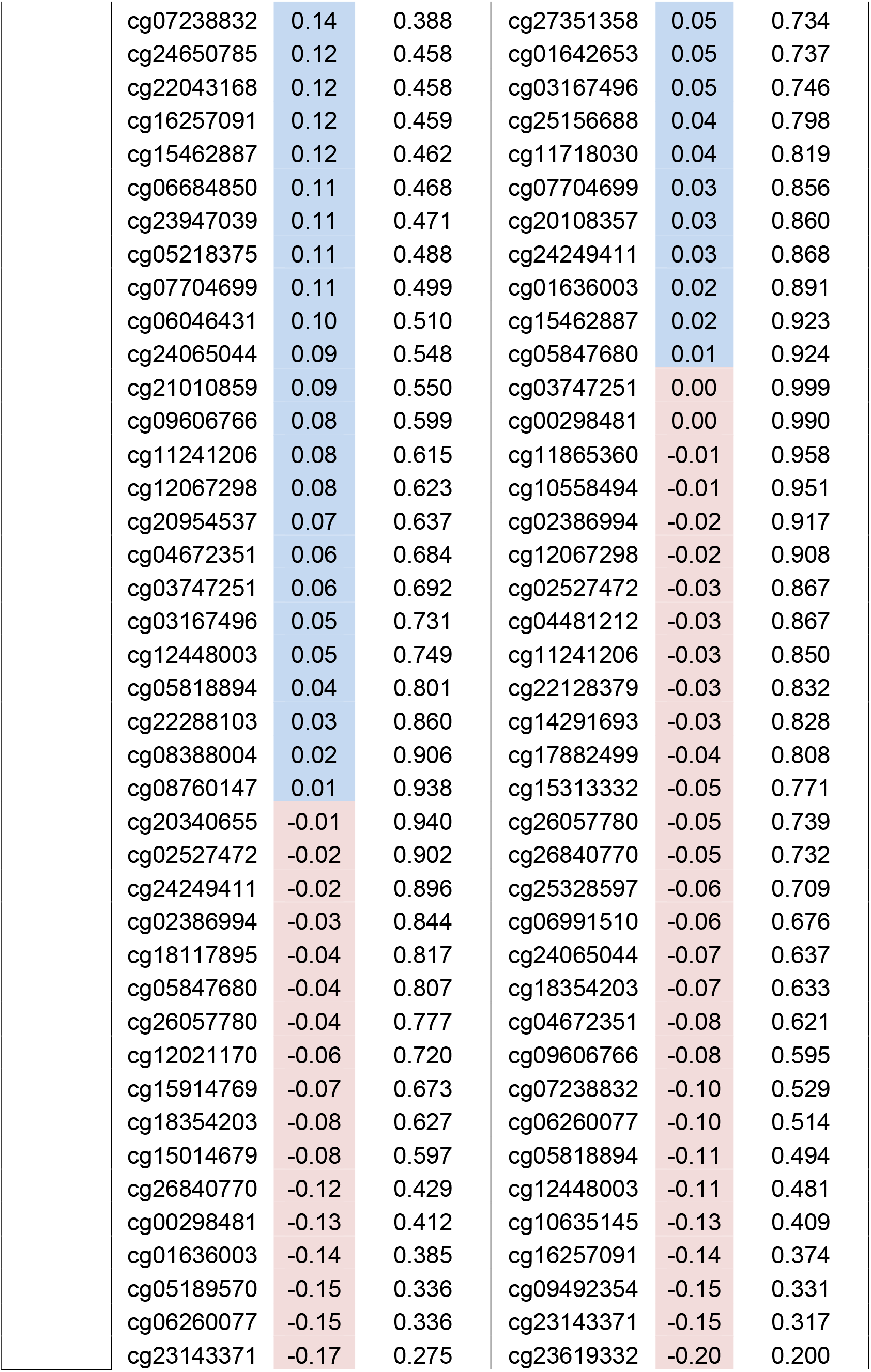
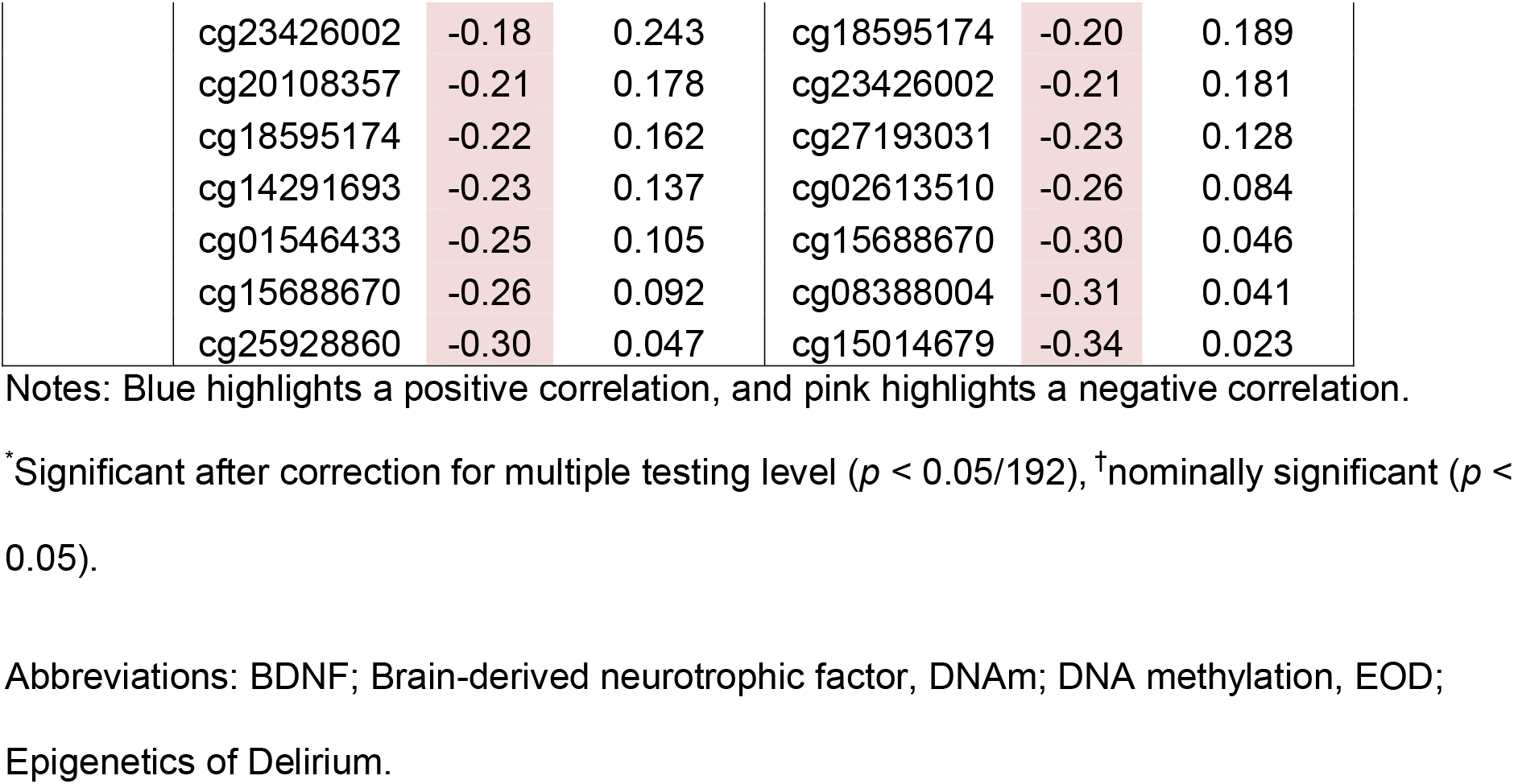
Correlation of age and blood DNAm at 83 CpGs in the *BDNF* gene compared between delirium cases vs non-delirium controls in the EOD cohort

| Gene | Delirium Cases (N=43) |  |  | Non Delirium Controls (N=44) |  |  |
| --- | --- | --- | --- | --- | --- | --- |
|  | CpG | r | p-value | CpG | r | p-value |
| *BDNF* | cg06816235<sup>*</sup> | 0.52 | 3.68E-04 | cg20340655<sup>†</sup> | 0.38 | 0.010 |
|  | cg05733135<sup>†</sup> | 0.43 | 0.004 | cg26949694<sup>†</sup> | 0.30 | 0.047 |
|  | cg13974632<sup>†</sup> | 0.39 | 0.010 | cg06046431<sup>†</sup> | 0.30 | 0.048 |
|  | cg25328597<sup>†</sup> | 0.37 | 0.015 | cg04106006 | 0.26 | 0.088 |
|  | cg10022526<sup>†</sup> | 0.36 | 0.017 | cg06816235 | 0.26 | 0.091 |
|  | cg09505801 | 0.30 | 0.052 | cg18117895 | 0.25 | 0.098 |
|  | cg11718030 | 0.30 | 0.053 | cg13974632 | 0.22 | 0.143 |
|  | cg01642653 | 0.29 | 0.060 | cg06684850 | 0.22 | 0.155 |
|  | cg27351358 | 0.26 | 0.087 | cg15710245 | 0.21 | 0.173 |
|  | cg04106006 | 0.26 | 0.092 | cg25457956 | 0.20 | 0.186 |
|  | cg07159484 | 0.25 | 0.100 | cg09505801 | 0.20 | 0.193 |
|  | cg01583131 | 0.25 | 0.103 | cg10022526 | 0.20 | 0.204 |
|  | cg17882499 | 0.24 | 0.124 | cg23947039 | 0.19 | 0.207 |
|  | cg10558494 | 0.23 | 0.134 | cg17413943 | 0.19 | 0.221 |
|  | cg15710245 | 0.23 | 0.145 | cg22043168 | 0.17 | 0.258 |
|  | cg23619332 | 0.22 | 0.161 | cg05218375 | 0.17 | 0.264 |
|  | cg25156688 | 0.21 | 0.177 | cg25928860 | 0.16 | 0.304 |
|  | cg06991510 | 0.21 | 0.183 | cg01225698 | 0.15 | 0.334 |
|  | cg08362738 | 0.20 | 0.193 | cg21010859 | 0.15 | 0.345 |
|  | cg17413943 | 0.19 | 0.215 | cg08362738 | 0.13 | 0.414 |
|  | cg09492354 | 0.19 | 0.217 | cg05733135 | 0.12 | 0.438 |
|  | cg25457956 | 0.18 | 0.244 | cg06979684 | 0.12 | 0.454 |
|  | cg04481212 | 0.17 | 0.280 | cg01583131 | 0.11 | 0.480 |
|  | cg15313332 | 0.17 | 0.287 | cg05189570 | 0.10 | 0.503 |
|  | cg02613510 | 0.16 | 0.298 | cg11806762 | 0.09 | 0.570 |
|  | cg22128379 | 0.16 | 0.307 | cg07159484 | 0.09 | 0.575 |
|  | cg11865360 | 0.16 | 0.315 | cg24650785 | 0.07 | 0.645 |
|  | cg03984780 | 0.15 | 0.331 | cg20954537 | 0.07 | 0.648 |
|  | cg27193031 | 0.15 | 0.337 | cg01546433 | 0.07 | 0.652 |
|  | cg10635145 | 0.15 | 0.341 | cg15914769 | 0.07 | 0.655 |
|  | cg26949694 | 0.15 | 0.342 | cg08760147 | 0.06 | 0.676 |
|  | cg06979684 | 0.15 | 0.349 | cg12021170 | 0.06 | 0.682 |
|  | cg11806762 | 0.14 | 0.361 | cg22288103 | 0.06 | 0.682 |
|  | cg06025631 | 0.14 | 0.369 | cg03984780 | 0.06 | 0.719 |
|  | cg01225698 | 0.14 | 0.387 | cg06025631 | 0.05 | 0.731 |

|  |  |  |  |  |  |
| --- | --- | --- | --- | --- | --- |
| cg07238832 | 0.14 | 0.388 | cg27351358 | 0.05 | 0.734 |
| cg24650785 | 0.12 | 0.458 | cg01642653 | 0.05 | 0.737 |
| cg22043168 | 0.12 | 0.458 | cg03167496 | 0.05 | 0.746 |
| cg16257091 | 0.12 | 0.459 | cg25156688 | 0.04 | 0.798 |
| cg15462887 | 0.12 | 0.462 | cg11718030 | 0.04 | 0.819 |
| cg06684850 | 0.11 | 0.468 | cg07704699 | 0.03 | 0.856 |
| cg23947039 | 0.11 | 0.471 | cg20108357 | 0.03 | 0.860 |
| cg05218375 | 0.11 | 0.488 | cg24249411 | 0.03 | 0.868 |
| cg07704699 | 0.11 | 0.499 | cg01636003 | 0.02 | 0.891 |
| cg06046431 | 0.10 | 0.510 | cg15462887 | 0.02 | 0.923 |
| cg24065044 | 0.09 | 0.548 | cg05847680 | 0.01 | 0.924 |
| cg21010859 | 0.09 | 0.550 | cg03747251 | 0.00 | 0.999 |
| cg09606766 | 0.08 | 0.599 | cg00298481 | 0.00 | 0.990 |
| cg11241206 | 0.08 | 0.615 | cg11865360 | -0.01 | 0.958 |
| cg12067298 | 0.08 | 0.623 | cg10558494 | -0.01 | 0.951 |
| cg20954537 | 0.07 | 0.637 | cg02386994 | -0.02 | 0.917 |
| cg04672351 | 0.06 | 0.684 | cg12067298 | -0.02 | 0.908 |
| cg03747251 | 0.06 | 0.692 | cg02527472 | -0.03 | 0.867 |
| cg03167496 | 0.05 | 0.731 | cg04481212 | -0.03 | 0.867 |
| cg12448003 | 0.05 | 0.749 | cg11241206 | -0.03 | 0.850 |
| cg05818894 | 0.04 | 0.801 | cg22128379 | -0.03 | 0.832 |
| cg22288103 | 0.03 | 0.860 | cg14291693 | -0.03 | 0.828 |
| cg08388004 | 0.02 | 0.906 | cg17882499 | -0.04 | 0.808 |
| cg08760147 | 0.01 | 0.938 | cg15313332 | -0.05 | 0.771 |
| cg20340655 | -0.01 | 0.940 | cg26057780 | -0.05 | 0.739 |
| cg02527472 | -0.02 | 0.902 | cg26840770 | -0.05 | 0.732 |
| cg24249411 | -0.02 | 0.896 | cg25328597 | -0.06 | 0.709 |
| cg02386994 | -0.03 | 0.844 | cg06991510 | -0.06 | 0.676 |
| cg18117895 | -0.04 | 0.817 | cg24065044 | -0.07 | 0.637 |
| cg05847680 | -0.04 | 0.807 | cg18354203 | -0.07 | 0.633 |
| cg26057780 | -0.04 | 0.777 | cg04672351 | -0.08 | 0.621 |
| cg12021170 | -0.06 | 0.720 | cg09606766 | -0.08 | 0.595 |
| cg15914769 | -0.07 | 0.673 | cg07238832 | -0.10 | 0.529 |
| cg18354203 | -0.08 | 0.627 | cg06260077 | -0.10 | 0.514 |
| cg15014679 | -0.08 | 0.597 | cg05818894 | -0.11 | 0.494 |
| cg26840770 | -0.12 | 0.429 | cg12448003 | -0.11 | 0.481 |
| cg00298481 | -0.13 | 0.412 | cg10635145 | -0.13 | 0.409 |
| cg01636003 | -0.14 | 0.385 | cg16257091 | -0.14 | 0.374 |
| cg05189570 | -0.15 | 0.336 | cg09492354 | -0.15 | 0.331 |
| cg06260077 | -0.15 | 0.336 | cg23143371 | -0.15 | 0.317 |
| cg23143371 | -0.17 | 0.275 | cg23619332 | -0.20 | 0.200 |
| cg23426002 | -0.18 | 0.243 | cg18595174 | -0.20 | 0.189 |
| cg20108357 | -0.21 | 0.178 | cg23426002 | -0.21 | 0.181 |
| cg18595174 | -0.22 | 0.162 | cg27193031 | -0.23 | 0.128 |
| cg14291693 | -0.23 | 0.137 | cg02613510 | -0.26 | 0.084 |
| cg01546433 | -0.25 | 0.105 | cg15688670 | -0.30 | 0.046 |
| cg15688670 | -0.26 | 0.092 | cg08388004 | -0.31 | 0.041 |
| cg25928860 | -0.30 | 0.047 | cg15014679 | -0.34 | 0.023 |

Notes: Blue highlights a positive correlation, and pink highlights a negative correlation.
\*Significant after correction for multiple testing level ( $p < 0.05/192$ ), <sup>†</sup>nominally significant ( $p < 0.05$ ).
Abbreviations: BDNF; Brain-derived neurotrophic factor, DNAm; DNA methylation, EOD; Epigenetics of Delirium.

## DISCUSSION

In the present study, DNAm of the neurotrophic genes were positively associated with aging in both the blood and brain samples. Furthermore, these DNAm were more positively correlated with aging in delirious patients than controls. These results support our hypothesis that DNAm in neurotrophic genes in both brain and blood increases with aging, and such changes are more significant among delirium patients.

Data from the GTP cohort blood samples showed that the majority (96 among 109 CpGs; 88.1%) of the nominally significant CpGs were positively correlated with aging. With the NSG cohort brain samples, we also found that the majority (55 among 62 CpGs; 88.7%) of the nominally significant CpGs were positively correlated with aging. Furthermore, the top hit CpG in the GTP study was also significantly positively correlated in the NSG study with brain samples. In total, 6 among the 16 significant CpGs after Bonferroni corrected genome-wide significance levels in the GTP cohort were also nominally significant in the NSG cohort. These results indicated that some neurotrophic CpGs show similar associations between aging and DNAm levels in brain and blood tissues.

In the EOD cohort more CpGs in neurotrophic genes were positively correlated with aging in delirium cases than in controls. This is consistent with our hypothesis that neurotrophic genes show greater DNAm changes with aging among elderly patients, indicating these genes may be dysregulated with altered expression levels compared to controls.

Studies of the relationship between circulating BDNF levels and delirium reveal mixed results. Higher plasma BDNF levels were associated with emergence of agitation in elderly patients after gastrointestinal surgery [45]. In patients with spine surgery, an intraoperative decline in plasma BDNF was greater in patients who developed delirium [46]. A study of ICU patients showed that serum level of *BDNF* was significantly higher in delirium patients than in non-delirious controls [47]. No significant differences between delirium cases and controls were found in both a study of oncology inpatients [48], nor in acute medical inpatients [49].

There are limitations to this study. First, the NSG cohort was limited in sample size due to the infrequent occurrence of neurosurgical cases. Second, both blood and brain tissues consist of heterogeneous cellular populations. Third, a peripheral tissue, blood, had to be used to assess DNAm levels in the EOD cohort of delirium patients despite brain being the main organ of interest for delirium. Fourth, longitudinal studies are needed to distinguish whether the neurotrophic genes show more significantly correlated DNAm levels with aging prior to delirium onset.

To the best of our knowledge, this is the first study to analyze the degree of correlation between DNAm in neurotrophic genes and age in delirium patients. We found 16 CpGs in *BDNF, GDNF, NR4A2, NPAS4, NR4A1*, and *ARC* to be significantly correlated with aging in the large GTP cohort with blood samples, 6 of these CpGs were also nominally significant in the NSG cohort of brain samples. Within the delirium cohort itself, 8 more CpGs in the neurotrophic genes were associated with aging in cases than controls with the majority showing positive correlation. These findings provide initial evidence that the neurotrophic genes may be epigenetically modulated with aging, and in patients with delirium, this process may be dysregulated.

