## Supplementary Tables for "DNA methylation change in neurotrophic genes with aging and delirium evidenced from three independent cohorts"

Supplementary Table 1 : Correlations between age and DNAm levels at 226 CpGs in neurotrophic genes in blood samples obtained from the GTP cohort

| Gene | CpG | t | R square | p |
| --- | --- | --- | --- | --- |
| *GDNF* | cg02328239^**^ | 9.86 | 0.22 | 1.54E-20 |
| *BDNF* | cg05733135^**^ | 8.18 | 0.17 | 4.49E-15 |
| *GDNF* | cg00049047^**^ | 7.43 | 0.12 | 7.74E-13 |
| *BDNF* | cg13974632^**^ | 7.38 | 0.14 | 1.06E-12 |
| *BDNF* | cg06816235^**^ | 7.29 | 0.13 | 1.88E-12 |
| *BDNF* | cg22043168^**^ | 7.07 | 0.12 | 7.92E-12 |
| *NR4A2* | cg11358945^**^ | 7.04 | 0.14 | 9.50E-12 |
| *NPAS4* | cg14699728^**^ | 6.55 | 0.12 | 1.86E-10 |
| *GDNF* | cg07442479^**^ | 6.55 | 0.12 | 1.88E-10 |
| *BDNF* | cg23947039^**^ | 6.22 | 0.10 | 1.33E-09 |
| *BDNF* | cg26949694^**^ | 6.15 | 0.09 | 1.95E-09 |
| *NR4A2* | cg00194126^**^ | 5.84 | 0.09 | 1.14E-08 |
| *NR4A1* | cg26446929^**^ | -5.74 | 0.07 | 1.92E-08 |
| *BDNF* | cg03167496^**^ | 5.71 | 0.10 | 2.25E-08 |
| *GDNF* | cg26789779^**^ | 5.66 | 0.09 | 2.99E-08 |
| *ARC* | cg16922810^**^ | -5.60 | 0.09 | 4.11E-08 |
| *BDNF* | cg01583131^*^ | 5.46 | 0.09 | 8.73E-08 |
| *BDNF* | cg25412831^*^ | 5.38 | 0.06 | 1.34E-07 |
| *NR4A2* | cg14811105^*^ | 5.36 | 0.08 | 1.47E-07 |
| *GDNF* | cg21590264^*^ | 5.30 | 0.08 | 1.94E-07 |
| *BDNF* | cg25962210^*^ | 5.18 | 0.09 | 3.58E-07 |
| *NR4A2* | cg02964555^*^ | 5.18 | 0.07 | 3.58E-07 |
| *NR4A2* | cg21226516^*^ | 5.14 | 0.08 | 4.47E-07 |
| *BDNF* | cg11718030^*^ | 4.97 | 0.07 | 1.01E-06 |
| *BDNF* | cg17413943^*^ | 4.90 | 0.05 | 1.41E-06 |
| *GDNF* | cg25602684^*^ | 4.88 | 0.08 | 1.59E-06 |
| *GDNF* | cg07715201^*^ | 4.79 | 0.06 | 2.40E-06 |
| *GDNF* | cg14492800^*^ | 4.77 | 0.08 | 2.62E-06 |
| *GDNF* | cg08204023^*^ | 4.76 | 0.07 | 2.73E-06 |
| *NPAS4* | cg04768203^*^ | 4.62 | 0.07 | 5.33E-06 |
| *GDNF* | cg26295057^*^ | 4.49 | 0.06 | 9.56E-06 |
| *GDNF* | cg18182111^*^ | 4.42 | 0.04 | 1.31E-05 |
| *BDNF* | cg04481212^*^ | 4.40 | 0.05 | 1.39E-05 |
| *BDNF* | cg21010859^*^ | 4.39 | 0.05 | 1.51E-05 |
| *GDNF* | cg21319053^*^ | 4.31 | 0.07 | 2.06E-05 |
| *BDNF* | cg24249411^*^ | 4.30 | 0.04 | 2.21E-05 |
| *FOS* | cg12061886^*^ | 4.18 | 0.03 | 3.64E-05 |
| *GDNF* | cg04209913^*^ | 4.18 | 0.04 | 3.66E-05 |
| *GDNF* | cg18725867^*^ | 4.17 | 0.06 | 3.78E-05 |
| *GDNF* | cg04442426^*^ | -4.11 | 0.07 | 4.82E-05 |
| *BDNF* | cg06684850^*^ | 4.09 | 0.05 | 5.29E-05 |
| *GDNF* | cg05330056^*^ | -4.07 | 0.19 | 5.73E-05 |
| *BDNF* | cg06991510^*^ | 4.05 | 0.04 | 6.11E-05 |
| *BDNF* | cg01225698^*^ | 4.04 | 0.03 | 6.57E-05 |
| *GDNF* | cg14590843^*^ | 3.98 | 0.03 | 8.41E-05 |
| *ARC* | cg19438565^*^ | 3.95 | 0.05 | 9.43E-05 |
| *NR4A2* | cg00426720^*^ | 3.93 | 0.04 | 1.03E-04 |
| *NR4A2* | cg10089963^*^ | 3.90 | 0.04 | 1.14E-04 |
| *GDNF* | cg07423205^*^ | 3.85 | 0.04 | 1.40E-04 |
| *BDNF* | cg25328597^*^ | 3.84 | 0.05 | 1.47E-04 |
| *BDNF* | cg18867480^*^ | 3.83 | 0.05 | 1.52E-04 |
| *NR4A2* | cg20398418 | 3.75 | 0.04 | 2.02E-04 |
| *NR4A2* | cg13500877 | 3.75 | 0.04 | 2.05E-04 |
| *GDNF* | cg15368455 | 3.69 | 0.04 | 2.55E-04 |
| *NPAS4* | cg23484234 | 3.68 | 0.04 | 2.66E-04 |
| *NPAS4* | cg08715791 | 3.62 | 0.06 | 3.34E-04 |
| *BDNF* | cg01642653 | 3.54 | 0.05 | 4.48E-04 |
| *BDNF* | cg10022526 | 3.42 | 0.04 | 0.001 |
| *GDNF* | cg24813562 | 3.28 | 0.04 | 0.001 |
| *NPAS4* | cg22134325 | 3.22 | 0.05 | 0.001 |
| *GDNF* | cg23097534 | 3.20 | 0.03 | 0.001 |
| *NPAS4* | cg13215078 | 3.19 | 0.06 | 0.002 |
| *BDNF* | cg26057780 | 3.18 | 0.04 | 0.002 |
| *ARC* | cg02675353 | 3.16 | 0.03 | 0.002 |
| *NR4A2* | cg18941818 | 3.14 | 0.03 | 0.002 |
| *GDNF* | cg04439218 | 3.14 | 0.04 | 0.002 |
| *NR4A2* | cg20570611 | 3.12 | 0.03 | 0.002 |
| *FOS* | cg15337055 | -3.10 | 0.02 | 0.002 |
| *BDNF* | cg08362738 | 2.98 | 0.03 | 0.003 |
| *GDNF* | cg07309340 | -2.98 | 0.13 | 0.003 |
| *NR4A1* | cg26933107 | -2.97 | 0.02 | 0.003 |
| *BDNF* | cg15462887 | 2.94 | 0.03 | 0.004 |
| *GDNF* | cg16408014 | -2.93 | 0.09 | 0.004 |
| *FOS* | cg14154718 | -2.92 | 0.03 | 0.004 |
| *BDNF* | cg10558494 | 2.88 | 0.01 | 0.004 |
| *NR4A2* | cg00240195 | 2.87 | 0.02 | 0.004 |
| *BDNF* | cg27351358 | 2.81 | 0.05 | 0.005 |
| *NR4A2* | cg09408520 | -2.69 | 0.01 | 0.007 |
| *BDNF* | cg25457956 | 2.69 | 0.01 | 0.007 |
| *BDNF* | cg14291693 | -2.68 | 0.03 | 0.008 |
| *NPAS4* | cg09835239 | 2.66 | 0.02 | 0.008 |
| *NR4A1* | cg13707690 | 2.64 | 0.01 | 0.009 |
| *FOS* | cg23404711 | 2.61 | 0.03 | 0.009 |
| *BDNF* | cg24650785 | 2.61 | 0.04 | 0.009 |
| *BDNF* | cg02527472 | 2.61 | 0.02 | 0.010 |
| *NR4A2* | cg00558219 | 2.59 | 0.01 | 0.010 |
| *BDNF* | cg03984780 | 2.57 | 0.01 | 0.011 |
| *NR4A2* | cg16058600 | 2.57 | 0.01 | 0.011 |
| *GDNF* | cg18237607 | 2.51 | 0.04 | 0.012 |
| *BDNF* | cg01418645 | 2.48 | 0.00 | 0.014 |
| *GDNF* | cg08307469 | 2.34 | 0.04 | 0.020 |
| *BDNF* | cg20340655 | 2.30 | 0.03 | 0.022 |
| *BDNF* | cg11241206 | 2.29 | 0.03 | 0.023 |
| *FOS* | cg20901874 | -2.24 | 0.00 | 0.025 |
| *ARC* | cg01509843 | 2.23 | 0.01 | 0.026 |
| *BDNF* | cg15313332 | 2.20 | 0.02 | 0.029 |
| *BDNF* | cg15710245 | 2.19 | 0.02 | 0.029 |
| *NR4A2* | cg03339537 | 2.19 | 0.01 | 0.029 |
| *NR4A2* | cg27074041 | 2.19 | 0.02 | 0.029 |
| *BDNF* | cg08388004 | 2.12 | 0.04 | 0.034 |
| *BDNF* | cg05818894 | 2.11 | 0.04 | 0.036 |
| *ARC* | cg24981018 | 2.10 | 0.04 | 0.037 |
| *BDNF* | cg16257091 | 2.08 | 0.00 | 0.039 |
| *NR4A1* | cg21368094 | -2.07 | 0.03 | 0.040 |
| *GDNF* | cg18121355 | 2.05 | 0.02 | 0.041 |
| *FOS* | cg16701133 | 2.03 | 0.00 | 0.043 |
| *NR4A2* | cg18881247 | 2.02 | 0.01 | 0.044 |
| *BDNF* | cg12448003 | 2.00 | 0.00 | 0.046 |
| *NR4A2* | cg21758126 | 1.98 | 0.01 | 0.048 |
| *GDNF* | cg07486383 | 1.96 | 0.03 | 0.051 |
| *BDNF* | cg01636003 | -1.96 | 0.04 | 0.051 |
| *BDNF* | cg07704699 | -1.95 | 0.01 | 0.052 |
| *BDNF* | cg18117895 | 1.95 | 0.00 | 0.052 |
| *NPAS4* | cg15273822 | 1.91 | 0.01 | 0.057 |
| *NR4A2* | cg06101180 | 1.86 | 0.02 | 0.063 |
| *BDNF* | cg23497217 | 1.84 | 0.00 | 0.067 |
| *GDNF* | cg26559974 | -1.81 | 0.05 | 0.071 |
| *ARC* | cg14225847 | 1.79 | 0.01 | 0.075 |
| *GDNF* | cg16138150 | 1.78 | 0.03 | 0.076 |
| *NR4A2* | cg23474904 | -1.76 | -0.01 | 0.079 |
| *BDNF* | cg00298481 | -1.72 | 0.01 | 0.086 |
| *NR4A2* | cg17654050 | 1.69 | 0.01 | 0.091 |
| *GDNF* | cg20683765 | 1.69 | 0.04 | 0.092 |
| *BDNF* | cg24377657 | 1.67 | 0.00 | 0.095 |
| *NR4A1* | cg04678231 | -1.65 | 0.00 | 0.099 |
| *BDNF* | cg23619332 | 1.57 | 0.03 | 0.117 |
| *ARC* | cg10104451 | -1.57 | -0.01 | 0.118 |
| *NR4A1* | cg03733196 | -1.55 | 0.02 | 0.123 |
| *NR4A2* | cg16151636 | -1.51 | -0.01 | 0.132 |
| *NR4A1* | cg03567117 | -1.50 | 0.01 | 0.135 |
| *NR4A2* | cg09906382 | 1.50 | 0.00 | 0.135 |
| *FOS* | cg01208563 | -1.46 | 0.00 | 0.144 |
| *BDNF* | cg06046431 | 1.36 | -0.01 | 0.175 |
| *BDNF* | cg09606766 | -1.34 | 0.00 | 0.180 |
| *BDNF* | cg26840770 | 1.34 | -0.02 | 0.182 |
| *BDNF* | cg05189570 | -1.33 | 0.02 | 0.185 |
| *GDNF* | cg17383727 | 1.29 | 0.04 | 0.198 |
| *BDNF* | cg20108357 | 1.27 | 0.01 | 0.204 |
| *GDNF* | cg00712841 | -1.27 | 0.08 | 0.204 |
| *GDNF* | cg02331025 | 1.27 | 0.03 | 0.206 |
| *FOS* | cg03509965 | 1.26 | 0.01 | 0.210 |
| *BDNF* | cg18354203 | 1.25 | 0.00 | 0.211 |
| *GDNF* | cg19622474 | 1.25 | 0.05 | 0.211 |
| *NR4A1* | cg06811166 | -1.24 | 0.01 | 0.216 |
| *ARC* | cg00487506 | -1.24 | 0.01 | 0.218 |
| *GDNF* | cg05462359 | -1.22 | 0.01 | 0.223 |
| *FOS* | cg18717355 | 1.22 | 0.01 | 0.225 |
| *NR4A1* | cg12541478 | 1.21 | -0.01 | 0.227 |
| *NR4A2* | cg07516970 | 1.20 | 0.00 | 0.231 |
| *NR4A2* | cg07646377 | -1.16 | -0.01 | 0.248 |
| *ARC* | cg05415840 | -1.15 | 0.01 | 0.250 |
| *BDNF* | cg20954537 | -1.15 | 0.01 | 0.250 |
| *NR4A2* | cg20804199 | -1.14 | -0.01 | 0.256 |
| *FOS* | cg10565512 | 1.11 | 0.00 | 0.268 |
| *NR4A2* | cg15699971 | -1.11 | 0.03 | 0.269 |
| *FOS* | cg10282345 | 1.09 | 0.02 | 0.277 |
| *BDNF* | cg02613510 | -1.07 | 0.00 | 0.287 |
| *GDNF* | cg02450613 | 1.04 | 0.01 | 0.298 |
| *NR4A2* | cg03953709 | -1.03 | 0.00 | 0.303 |
| *GDNF* | cg07295550 | -0.99 | 0.02 | 0.325 |
| *BDNF* | cg04672351 | 0.96 | 0.04 | 0.340 |
| *BDNF* | cg05218375 | 0.95 | 0.00 | 0.344 |
| *BDNF* | cg04106006 | -0.94 | 0.01 | 0.348 |
| *BDNF* | cg27193031 | 0.93 | 0.02 | 0.354 |
| *BDNF* | cg07159484 | 0.92 | 0.00 | 0.360 |
| *NR4A2* | cg13945301 | -0.91 | 0.00 | 0.361 |
| *FOS* | cg13819869 | 0.91 | -0.02 | 0.362 |
| *NPAS4* | cg11635197 | 0.88 | 0.00 | 0.378 |
| *ARC* | cg13172906 | -0.88 | 0.03 | 0.380 |
| *GDNF* | cg07128111 | 0.87 | 0.05 | 0.383 |
| *NPAS4* | cg02055483 | 0.84 | 0.00 | 0.401 |
| *BDNF* | cg25381667 | -0.84 | 0.00 | 0.403 |
| *GDNF* | cg01107142 | 0.83 | -0.01 | 0.407 |
| *BDNF* | cg15914769 | -0.83 | 0.02 | 0.408 |
| *BDNF* | cg23426002 | -0.81 | 0.02 | 0.418 |
| *GDNF* | cg23400942 | -0.77 | 0.05 | 0.443 |
| *NR4A2* | cg16246410 | -0.75 | 0.01 | 0.451 |
| *FOS* | cg17644867 | 0.75 | -0.01 | 0.451 |
| *BDNF* | cg15014679 | -0.75 | 0.03 | 0.451 |
| *GDNF* | cg12930882 | 0.68 | 0.02 | 0.495 |
| *BDNF* | cg07238832 | 0.67 | 0.00 | 0.506 |
| *ARC* | cg08387463 | -0.63 | 0.01 | 0.527 |
| *NPAS4* | cg24435401 | 0.63 | -0.01 | 0.532 |
| *NR4A2* | cg20945253 | -0.62 | 0.00 | 0.534 |
| *NR4A1* | cg07544244 | 0.62 | -0.01 | 0.537 |
| *NR4A1* | cg12610867 | -0.61 | -0.01 | 0.543 |
| *BDNF* | cg10635145 | -0.60 | 0.03 | 0.550 |
| *BDNF* | cg11806762 | -0.58 | 0.01 | 0.560 |
| *FOS* | cg25975379 | -0.55 | -0.02 | 0.583 |
| *NR4A2* | cg24651504 | 0.54 | 0.01 | 0.586 |
| *FOS* | cg25034766 | 0.54 | 0.02 | 0.590 |
| *BDNF* | cg09492354 | 0.53 | -0.01 | 0.597 |
| *BDNF* | cg24065044 | 0.53 | 0.00 | 0.598 |
| *FOS* | cg00773696 | -0.51 | 0.00 | 0.611 |
| *GDNF* | cg05730365 | 0.51 | 0.02 | 0.613 |
| *NR4A1* | cg20718727 | -0.50 | -0.01 | 0.615 |
| *ARC* | cg23210049 | -0.50 | 0.03 | 0.616 |
| *NPAS4* | cg23667955 | 0.49 | 0.02 | 0.622 |
| *ARC* | cg04321580 | -0.47 | -0.01 | 0.637 |
| *BDNF* | cg06025631 | -0.46 | 0.00 | 0.643 |
| *FOS* | cg11872076 | -0.45 | 0.00 | 0.651 |
| *GDNF* | cg26077283 | 0.45 | 0.00 | 0.653 |
| *ARC* | cg01577292 | -0.44 | -0.02 | 0.657 |
| *NR4A1* | cg18713088 | -0.44 | 0.00 | 0.659 |
| *BDNF* | cg14589148 | 0.39 | -0.01 | 0.698 |
| *BDNF* | cg15688670 | 0.37 | 0.02 | 0.710 |
| *NPAS4* | cg14326225 | 0.37 | 0.03 | 0.710 |
| *NR4A1* | cg05486650 | -0.36 | 0.00 | 0.720 |
| *NPAS4* | cg19568845 | 0.35 | 0.00 | 0.725 |
| *GDNF* | cg01043865 | 0.33 | 0.00 | 0.739 |
| *BDNF* | cg06260077 | 0.32 | 0.00 | 0.746 |
| *BDNF* | cg03747251 | -0.32 | -0.02 | 0.751 |
| *ARC* | cg24450303 | 0.28 | 0.02 | 0.783 |
| *NPAS4* | cg25233308 | -0.27 | -0.01 | 0.791 |
| *NR4A2* | cg25247969 | -0.25 | -0.01 | 0.799 |
| *NR4A2* | cg11379337 | -0.23 | -0.01 | 0.816 |
| *GDNF* | cg26473844 | -0.23 | 0.01 | 0.821 |
| *BDNF* | cg18595174 | -0.22 | 0.00 | 0.824 |
| *FOS* | cg14102251 | -0.21 | -0.01 | 0.832 |
| *BDNF* | cg06979684 | 0.21 | -0.01 | 0.834 |
| *GDNF* | cg11275487 | -0.18 | 0.04 | 0.856 |
| *FOS* | cg07159858 | -0.17 | -0.01 | 0.864 |
| *GDNF* | cg21918513 | 0.15 | -0.01 | 0.881 |
| *NR4A2* | cg01123282 | -0.13 | -0.01 | 0.894 |
| *ARC* | cg08860119 | -0.11 | -0.01 | 0.914 |
| *GDNF* | cg20380689 | 0.02 | -0.02 | 0.982 |

Notes: Blue highlights a positive correlation, and pink highlights a negative correlation. ^**^Significant after Bonferroni corrected genome-wide significance levels (p < 5 × E-8), ^*^Significant after correction for multiple testing level (*p* < 0.05/226).

Abbreviations: AP-1 Transcription Factor Subunit, ARC; activity regulated cytoskeleton associated protein, BDNF; Brain-derived neurotrophic factor, FOS; Fos Proto-Oncogene, GDNF; glial cell-derived neurotrophic factor, GTP; Grady Trauma Project, NPAS4; neuronal Per-Arnt-Sim domain protein 4, NR4A1; nuclear receptor subfamily 4A1, NR4A2; nuclear receptor subfamily 4A2.

Supplementary Table 2 : Correlations between age and DNAm levels at 201 CpGs in neurotrophic genes in brain samples obtained from the NSG cohort

| cgid | Chr | Gene | Rho | p |
| --- | --- | --- | --- | --- |
| cg08715791^*^ | 11 | *NPAS4* | 0.85 | 1.17E-06 |
| cg02328239^*^ | 5 | *GDNF* | 0.85 | 1.26E-06 |
| cg27074041^*^ | 2 | *NR4A2* | 0.82 | 4.87E-06 |
| cg00558219^*^ | 2 | *NR4A2* | 0.75 | 9.09E-05 |
| cg26949694^*^ | 11 | *BDNF* | 0.75 | 9.49E-05 |
| cg03230469^*^ | 5 | *GDNF* | 0.73 | 1.67E-04 |
| cg02964555^†^ | 2 | *NR4A2* | 0.70 | 3.85E-04 |
| cg12296752^†^ | 11 | *BDNF* | 0.70 | 4.28E-04 |
| cg04407962^†^ | 5 | *GDNF* | 0.69 | 5.65E-04 |
| cg20570611^†^ | 2 | *NR4A2* | 0.68 | 6.67E-04 |
| cg04768203^†^ | 11 | *NPAS4* | 0.67 | 9.36E-04 |
| cg12930882^†^ | 5 | *GDNF* | 0.64 | 0.002 |
| cg20108357^†^ | 11 | *BDNF* | 0.63 | 0.002 |
| cg04209913^†^ | 5 | *GDNF* | 0.62 | 0.003 |
| cg14811105^†^ | 2 | *NR4A2* | 0.62 | 0.003 |
| cg07238832^†^ | 11 | *BDNF* | 0.61 | 0.003 |
| cg06684850^†^ | 11 | *BDNF* | 0.60 | 0.004 |
| cg11358945^†^ | 2 | *NR4A2* | 0.60 | 0.004 |
| cg21286419^†^ | 5 | *GDNF* | 0.60 | 0.004 |
| cg16058600^†^ | 2 | *NR4A2* | 0.58 | 0.006 |
| cg11209121^†^ | 2 | *NR4A2* | 0.58 | 0.006 |
| cg01107142^†^ | 5 | *GDNF* | 0.57 | 0.007 |
| cg26559974^†^ | 5 | *GDNF* | 0.57 | 0.007 |
| cg01123282^†^ | 2 | *NR4A2* | -0.57 | 0.007 |
| cg04106006^†^ | 11 | *BDNF* | 0.57 | 0.007 |
| cg00194126^†^ | 2 | *NR4A2* | 0.56 | 0.008 |
| cg17654050^†^ | 2 | *NR4A2* | 0.56 | 0.008 |
| cg15368455^†^ | 5 | *GDNF* | 0.56 | 0.008 |
| cg14291693^†^ | 11 | *BDNF* | 0.56 | 0.009 |
| cg14590843^†^ | 5 | *GDNF* | 0.56 | 0.009 |
| cg08760147^†^ | 11 | *BDNF* | 0.55 | 0.010 |
| cg00761985^†^ | 5 | *GDNF* | 0.55 | 0.010 |
| cg14617996^†^ | 2 | *NR4A2* | 0.54 | 0.011 |
| cg18595174^†^ | 11 | *BDNF* | 0.54 | 0.012 |
| cg05189570^†^ | 11 | *BDNF* | 0.53 | 0.013 |
| cg15710245^†^ | 11 | *BDNF* | -0.53 | 0.013 |
| cg13500877^†^ | 2 | *NR4A2* | 0.53 | 0.014 |
| cg01583131^†^ | 11 | *BDNF* | 0.53 | 0.014 |
| cg00240195^†^ | 2 | *NR4A2* | 0.52 | 0.015 |
| cg25928860^†^ | 11 | *BDNF* | 0.52 | 0.016 |
| cg10558494^†^ | 11 | *BDNF* | -0.52 | 0.016 |
| cg03167496^†^ | 11 | *BDNF* | 0.51 | 0.018 |
| cg23097534^†^ | 5 | *GDNF* | 0.51 | 0.018 |
| cg06816235^†^ | 11 | *BDNF* | 0.51 | 0.019 |
| cg06101180^†^ | 2 | *NR4A2* | 0.51 | 0.019 |
| cg06260077^†^ | 11 | *BDNF* | 0.50 | 0.021 |
| cg16246410^†^ | 2 | *NR4A2* | -0.50 | 0.021 |
| cg17413943^†^ | 11 | *BDNF* | 0.49 | 0.024 |
| cg11932911^†^ | 2 | *NR4A2* | 0.49 | 0.024 |
| cg11399967^†^ | 2 | *NR4A2* | 0.49 | 0.024 |
| cg09606766^†^ | 11 | *BDNF* | -0.49 | 0.025 |
| cg10089963^†^ | 2 | *NR4A2* | 0.49 | 0.025 |
| cg21226516^†^ | 2 | *NR4A2* | 0.48 | 0.027 |
| cg02331025^†^ | 5 | *GDNF* | 0.48 | 0.027 |
| cg23400942^†^ | 5 | *GDNF* | 0.47 | 0.030 |
| cg19717018^†^ | 5 | *GDNF* | 0.47 | 0.031 |
| cg10635145^†^ | 11 | *BDNF* | 0.46 | 0.035 |
| cg15699971^†^ | 2 | *NR4A2* | -0.46 | 0.036 |
| cg07159484^†^ | 11 | *BDNF* | -0.44 | 0.044 |
| cg03339537^†^ | 2 | *NR4A2* | 0.44 | 0.045 |
| cg18941818^†^ | 2 | *NR4A2* | 0.44 | 0.048 |
| cg01546433^†^ | 11 | *BDNF* | 0.43 | 0.049 |
| cg21506655 | 2 | *NR4A2* | 0.43 | 0.050 |
| cg01642653 | 11 | *BDNF* | 0.43 | 0.051 |
| cg07295550 | 5 | *GDNF* | -0.43 | 0.053 |
| cg21758126 | 2 | *NR4A2* | 0.43 | 0.054 |
| cg20420783 | 11 | *NPAS4* | 0.42 | 0.059 |
| cg12448003 | 11 | *BDNF* | -0.42 | 0.061 |
| cg24377657 | 11 | *BDNF* | -0.42 | 0.061 |
| cg23426002 | 11 | *BDNF* | 0.41 | 0.062 |
| cg04672351 | 11 | *BDNF* | 0.41 | 0.064 |
| cg12021170 | 11 | *BDNF* | 0.40 | 0.069 |
| cg17075252 | 11 | *BDNF* | 0.40 | 0.070 |
| cg18881247 | 2 | *NR4A2* | -0.40 | 0.074 |
| cg04442426 | 5 | *GDNF* | -0.40 | 0.075 |
| cg11275487 | 5 | *GDNF* | 0.40 | 0.075 |
| cg12067298 | 11 | *BDNF* | 0.40 | 0.076 |
| cg16815025 | 5 | *GDNF* | -0.39 | 0.079 |
| cg09492354 | 11 | *BDNF* | -0.39 | 0.080 |
| cg26473844 | 5 | *GDNF* | -0.39 | 0.084 |
| cg11865360 | 11 | *BDNF* | 0.38 | 0.085 |
| cg09906382 | 2 | *NR4A2* | 0.38 | 0.085 |
| cg25127975 | 2 | *NR4A2* | 0.37 | 0.103 |
| cg19622474 | 5 | *GDNF* | 0.36 | 0.105 |
| cg20683765 | 5 | *GDNF* | 0.36 | 0.109 |
| cg05730365 | 5 | *GDNF* | -0.36 | 0.113 |
| cg17882499 | 11 | *BDNF* | -0.35 | 0.117 |
| cg20804199 | 2 | *NR4A2* | -0.35 | 0.117 |
| cg05330056 | 5 | *GDNF* | -0.35 | 0.119 |
| cg06979684 | 11 | *BDNF* | 0.34 | 0.126 |
| cg16257091 | 11 | *BDNF* | 0.34 | 0.133 |
| cg16198719 | 5 | *GDNF* | -0.34 | 0.136 |
| cg17383727 | 5 | *GDNF* | 0.33 | 0.138 |
| cg16408014 | 5 | *GDNF* | -0.33 | 0.139 |
| cg26077283 | 5 | *GDNF* | -0.33 | 0.142 |
| cg05462359 | 5 | *GDNF* | -0.33 | 0.146 |
| cg00712841 | 5 | *GDNF* | 0.33 | 0.146 |
| cg01636003 | 11 | *BDNF* | -0.32 | 0.152 |
| cg23667955 | 11 | *NPAS4* | 0.32 | 0.153 |
| cg15914769 | 11 | *BDNF* | -0.32 | 0.154 |
| cg07128111 | 5 | *GDNF* | 0.32 | 0.154 |
| cg11718030 | 11 | *BDNF* | 0.32 | 0.156 |
| cg24435401 | 11 | *NPAS4* | 0.32 | 0.160 |
| cg20340655 | 11 | *BDNF* | -0.32 | 0.161 |
| cg06046431 | 11 | *BDNF* | -0.31 | 0.168 |
| cg18117895 | 11 | *BDNF* | 0.31 | 0.169 |
| cg11396695 | 5 | *GDNF* | -0.31 | 0.172 |
| cg02386994 | 11 | *BDNF* | 0.31 | 0.174 |
| cg14492800 | 5 | *GDNF* | -0.30 | 0.183 |
| cg02450613 | 5 | *GDNF* | -0.30 | 0.185 |
| cg22134325 | 11 | *NPAS4* | 0.30 | 0.192 |
| cg25156688 | 11 | *BDNF* | 0.29 | 0.194 |
| cg26295057 | 5 | *GDNF* | -0.29 | 0.195 |
| cg22043168 | 11 | *BDNF* | 0.29 | 0.198 |
| cg21010859 | 11 | *BDNF* | -0.29 | 0.203 |
| cg11806762 | 11 | *BDNF* | 0.29 | 0.204 |
| cg20398418 | 2 | *NR4A2* | 0.29 | 0.205 |
| cg07423205 | 5 | *GDNF* | -0.28 | 0.212 |
| cg13974632 | 11 | *BDNF* | 0.27 | 0.237 |
| cg24650785 | 11 | *BDNF* | 0.27 | 0.240 |
| cg25247969 | 2 | *NR4A2* | -0.26 | 0.256 |
| cg09505801 | 11 | *BDNF* | -0.26 | 0.258 |
| cg15688670 | 11 | *BDNF* | -0.26 | 0.259 |
| cg03612055 | 5 | *GDNF* | -0.25 | 0.275 |
| cg08204023 | 5 | *GDNF* | 0.25 | 0.278 |
| cg23474904 | 2 | *NR4A2* | 0.24 | 0.288 |
| cg14699728 | 11 | *NPAS4* | -0.24 | 0.298 |
| cg06991510 | 11 | *BDNF* | -0.24 | 0.303 |
| cg08362738 | 11 | *BDNF* | 0.23 | 0.313 |
| cg05818894 | 11 | *BDNF* | 0.23 | 0.316 |
| cg05733135 | 11 | *BDNF* | -0.22 | 0.333 |
| cg03266646 | 5 | *GDNF* | -0.22 | 0.348 |
| cg27351358 | 11 | *BDNF* | -0.21 | 0.352 |
| cg10022526 | 11 | *BDNF* | 0.20 | 0.384 |
| cg09835239 | 11 | *NPAS4* | -0.20 | 0.386 |
| cg25233308 | 11 | *NPAS4* | -0.19 | 0.402 |
| cg19568845 | 11 | *NPAS4* | 0.19 | 0.417 |
| cg07486383 | 5 | *GDNF* | -0.18 | 0.430 |
| cg07108936 | 2 | *NR4A2* | -0.18 | 0.430 |
| cg23947039 | 11 | *BDNF* | 0.18 | 0.437 |
| cg20945253 | 2 | *NR4A2* | -0.18 | 0.447 |
| cg02613510 | 11 | *BDNF* | 0.17 | 0.449 |
| cg22288103 | 11 | *BDNF* | -0.16 | 0.500 |
| cg02055483 | 11 | *NPAS4* | 0.15 | 0.507 |
| cg16151636 | 2 | *NR4A2* | -0.15 | 0.509 |
| cg23143371 | 11 | *BDNF* | 0.15 | 0.512 |
| cg11379337 | 2 | *NR4A2* | -0.15 | 0.525 |
| cg24249411 | 11 | *BDNF* | 0.14 | 0.533 |
| cg26789779 | 5 | *GDNF* | -0.14 | 0.544 |
| cg21918513 | 5 | *GDNF* | 0.14 | 0.555 |
| cg00049047 | 5 | *GDNF* | 0.13 | 0.565 |
| cg03953709 | 2 | *NR4A2* | -0.13 | 0.567 |
| cg03984780 | 11 | *BDNF* | 0.13 | 0.576 |
| cg11241206 | 11 | *BDNF* | -0.13 | 0.580 |
| cg13945301 | 2 | *NR4A2* | -0.13 | 0.580 |
| cg11635197 | 11 | *NPAS4* | -0.13 | 0.588 |
| cg04481212 | 11 | *BDNF* | 0.12 | 0.600 |
| cg07704699 | 11 | *BDNF* | 0.12 | 0.602 |
| cg03747251 | 11 | *BDNF* | -0.12 | 0.602 |
| cg09408520 | 2 | *NR4A2* | -0.12 | 0.604 |
| cg26840770 | 11 | *BDNF* | -0.12 | 0.608 |
| cg18237607 | 5 | *GDNF* | -0.11 | 0.630 |
| cg25328597 | 11 | *BDNF* | -0.11 | 0.648 |
| cg23484234 | 11 | *NPAS4* | 0.11 | 0.648 |
| cg18354203 | 11 | *BDNF* | 0.10 | 0.652 |
| cg10087138 | 11 | *NPAS4* | 0.10 | 0.680 |
| cg25381667 | 11 | *BDNF* | 0.09 | 0.687 |
| cg22128379 | 11 | *BDNF* | 0.09 | 0.691 |
| cg05218375 | 11 | *BDNF* | -0.08 | 0.716 |
| cg25457956 | 11 | *BDNF* | -0.08 | 0.722 |
| cg24398268 | 5 | *GDNF* | -0.08 | 0.731 |
| cg25412831 | 11 | *BDNF* | 0.08 | 0.733 |
| cg24065044 | 11 | *BDNF* | -0.07 | 0.754 |
| cg26057780 | 11 | *BDNF* | -0.07 | 0.754 |
| cg07646377 | 2 | *NR4A2* | -0.07 | 0.765 |
| cg15462887 | 11 | *BDNF* | 0.07 | 0.771 |
| cg18725867 | 5 | *GDNF* | -0.06 | 0.797 |
| cg01225698 | 11 | *BDNF* | -0.06 | 0.806 |
| cg04439218 | 5 | *GDNF* | -0.06 | 0.806 |
| cg05847680 | 11 | *BDNF* | -0.05 | 0.816 |
| cg16138150 | 5 | *GDNF* | -0.05 | 0.821 |
| cg07715201 | 5 | *GDNF* | 0.05 | 0.823 |
| cg21590264 | 5 | *GDNF* | -0.05 | 0.836 |
| cg20954537 | 11 | *BDNF* | -0.05 | 0.840 |
| cg25565192 | 11 | *NPAS4* | 0.04 | 0.851 |
| cg07309340 | 5 | *GDNF* | -0.04 | 0.862 |
| cg13215078 | 11 | *NPAS4* | 0.04 | 0.865 |
| cg15273822 | 11 | *NPAS4* | 0.04 | 0.871 |
| cg00298481 | 11 | *BDNF* | -0.03 | 0.882 |
| cg08388004 | 11 | *BDNF* | -0.03 | 0.895 |
| cg15313332 | 11 | *BDNF* | 0.03 | 0.898 |
| cg14589148 | 11 | *BDNF* | -0.02 | 0.918 |
| cg15014679 | 11 | *BDNF* | -0.02 | 0.924 |
| cg25602684 | 5 | *GDNF* | 0.02 | 0.924 |
| cg23619332 | 11 | *BDNF* | 0.02 | 0.931 |
| cg07442479 | 5 | *GDNF* | -0.02 | 0.931 |
| cg24533810 | 5 | *GDNF* | -0.02 | 0.942 |
| cg14326225 | 11 | *NPAS4* | -0.02 | 0.947 |
| cg27193031 | 11 | *BDNF* | -0.01 | 0.953 |
| cg06025631 | 11 | *BDNF* | 0.01 | 0.955 |
| cg02527472 | 11 | *BDNF* | -0.01 | 0.973 |

Notes: Blue highlights a positive correlation, and pink highlights a negative correlation. ^*^Significant after correction for multiple testing level (*p* < 0.05/201), ^†^nominally significant (*p* < 0.05).

Abbreviations: BDNF; Brain-derived neurotrophic factor, GDNF; glial cell-derived neurotrophic factor, NPAS4; neuronal Per-Arnt-Sim domain protein 4, NR4A2; nuclear receptor subfamily 4A2, NSG; neurosurgery.

Supplementary Table 3 : Correlation of age and blood DNAm at 52 CpGs in the *GDNF* gene compared between delirium cases vs non-delirium controls in the EOD cohort

| Gene | Delirium Cases (N=43) | | | Non Delirium Controls (N=44) | | |
| --- | --- | --- | --- | --- | --- | --- |
|  | CpG | r | p-value | CpG | r | p-value |
| *GDNF* | cg26295057 | 0.37 | 0.014^*^ | cg07715201 | 0.42 | 0.004^*^ |
|  | cg03230469 | 0.34 | 0.024^*^ | cg14492800 | 0.30 | 0.045^*^ |
|  | cg07442479 | 0.32 | 0.035^*^ | cg16815025 | 0.29 | 0.060 |
|  | cg21286419 | 0.32 | 0.038^*^ | cg12930882 | 0.28 | 0.069 |
|  | cg02450613 | 0.31 | 0.042^*^ | cg19622474 | 0.26 | 0.089 |
|  | cg14492800 | 0.29 | 0.057 | cg19717018 | 0.24 | 0.115 |
|  | cg24533810 | 0.28 | 0.065 | cg04209913 | 0.24 | 0.121 |
|  | cg18725867 | 0.26 | 0.086 | cg16198719 | 0.22 | 0.160 |
|  | cg25602684 | 0.26 | 0.086 | cg07423205 | 0.17 | 0.270 |
|  | cg11396695 | 0.26 | 0.090 | cg07442479 | 0.17 | 0.279 |
|  | cg04407962 | 0.25 | 0.111 | cg24398268 | 0.17 | 0.279 |
|  | cg16198719 | 0.24 | 0.125 | cg02328239 | 0.17 | 0.284 |
|  | cg21590264 | 0.22 | 0.157 | cg07295550 | 0.11 | 0.469 |
|  | cg24398268 | 0.21 | 0.168 | cg24533810 | 0.10 | 0.537 |
|  | cg03612055 | 0.21 | 0.170 | cg03612055 | 0.09 | 0.556 |
|  | cg16138150 | 0.21 | 0.176 | cg03230469 | 0.09 | 0.559 |
|  | cg05730365 | 0.20 | 0.189 | cg00049047 | 0.09 | 0.582 |
|  | cg19717018 | 0.19 | 0.224 | cg07128111 | 0.08 | 0.621 |
|  | cg00049047 | 0.19 | 0.224 | cg18237607 | 0.07 | 0.658 |
|  | cg07715201 | 0.19 | 0.227 | cg26295057 | 0.07 | 0.667 |
|  | cg23097534 | 0.19 | 0.227 | cg03266646 | 0.06 | 0.678 |
|  | cg11275487 | 0.18 | 0.261 | cg05462359 | 0.06 | 0.684 |
|  | cg03266646 | 0.15 | 0.323 | cg25602684 | 0.06 | 0.700 |
|  | cg07423205 | 0.15 | 0.343 | cg16138150 | 0.06 | 0.712 |
|  | cg21918513 | 0.15 | 0.351 | cg08204023 | 0.05 | 0.768 |
|  | cg20683765 | 0.12 | 0.435 | cg18725867 | 0.04 | 0.777 |
|  | cg14590843 | 0.11 | 0.495 | cg21590264 | 0.04 | 0.802 |
|  | cg16408014 | 0.10 | 0.510 | cg01107142 | 0.04 | 0.814 |
|  | cg05462359 | 0.10 | 0.514 | cg02450613 | 0.03 | 0.864 |
|  | cg08204023 | 0.10 | 0.529 | cg15368455 | 0.03 | 0.870 |
|  | cg18237607 | 0.10 | 0.544 | cg26077283 | -0.04 | 0.803 |
|  | cg26077283 | 0.08 | 0.619 | cg14590843 | -0.05 | 0.762 |
|  | cg07128111 | 0.07 | 0.648 | cg04439218 | -0.07 | 0.666 |
|  | cg07295550 | 0.04 | 0.787 | cg00761985 | -0.10 | 0.536 |
|  | cg04209913 | 0.04 | 0.799 | cg20683765 | -0.10 | 0.527 |
|  | cg17383727 | 0.02 | 0.875 | cg21918513 | -0.12 | 0.427 |
|  | cg12930882 | 0.02 | 0.895 | cg16408014 | -0.13 | 0.401 |
|  | cg02328239 | 0.00 | 0.977 | cg07309340 | -0.13 | 0.388 |
|  | cg00712841 | 0.00 | 0.975 | cg11275487 | -0.14 | 0.376 |
|  | cg01107142 | -0.07 | 0.659 | cg07486383 | -0.14 | 0.372 |
|  | cg19622474 | -0.08 | 0.599 | cg23400942 | -0.14 | 0.360 |
|  | cg16815025 | -0.09 | 0.587 | cg23097534 | -0.14 | 0.356 |
|  | cg23400942 | -0.09 | 0.567 | cg02331025 | -0.15 | 0.342 |
|  | cg07486383 | -0.12 | 0.440 | cg17383727 | -0.16 | 0.302 |
|  | cg02331025 | -0.14 | 0.373 | cg05730365 | -0.16 | 0.294 |
|  | cg04439218 | -0.15 | 0.343 | cg00712841 | -0.17 | 0.275 |
|  | cg26559974 | -0.17 | 0.283 | cg11396695 | -0.17 | 0.272 |
|  | cg00761985 | -0.19 | 0.221 | cg26559974 | -0.18 | 0.230 |
|  | cg04442426 | -0.22 | 0.160 | cg05330056 | -0.24 | 0.114 |
|  | cg15368455 | -0.23 | 0.143 | cg21286419 | -0.27 | 0.076 |
|  | cg07309340 | -0.28 | 0.067 | cg04442426 | -0.37 | 0.012 |
|  | cg05330056 | -0.31 | 0.046 | cg04407962 | -0.40 | 0.007 |

Notes: Blue highlights a positive correlation, and pink highlights a negative correlation. ^*^Nominally significant (*p* < 0.05).

Abbreviations: GDNF; glial cell-derived neurotrophic factor, DNAm; DNA methylation, EOD; Epigenetics of Delirium.

Supplementary Table 4 : Correlation of age and blood DNAm at 39 CpGs in the *NR4A2* gene compared between delirium cases vs non-delirium controls in the EOD cohort

| Gene | Delirium Cases (N=43) | | | Non Delirium Controls (N=44) | | |
| --- | --- | --- | --- | --- | --- | --- |
|  | CpG | r | p-value | CpG | r | p-value |
| *NR4A2* | cg00558219 | 0.37 | 0.016^*^ | cg25247969 | 0.24 | 0.123 |
|  | cg10089963 | 0.32 | 0.038^*^ | cg07108936 | 0.21 | 0.176 |
|  | cg20804199 | 0.30 | 0.048^*^ | cg10089963 | 0.19 | 0.215 |
|  | cg21506655 | 0.27 | 0.080 | cg00558219 | 0.17 | 0.281 |
|  | cg13500877 | 0.25 | 0.112 | cg15699971 | 0.15 | 0.343 |
|  | cg02964555 | 0.24 | 0.128 | cg14811105 | 0.14 | 0.349 |
|  | cg21758126 | 0.22 | 0.165 | cg20570611 | 0.11 | 0.480 |
|  | cg13945301 | 0.19 | 0.230 | cg20398418 | 0.10 | 0.533 |
|  | cg03953709 | 0.18 | 0.248 | cg20945253 | 0.10 | 0.533 |
|  | cg09906382 | 0.14 | 0.357 | cg02964555 | 0.09 | 0.552 |
|  | cg00194126 | 0.11 | 0.496 | cg16058600 | 0.09 | 0.578 |
|  | cg23474904 | 0.10 | 0.507 | cg21758126 | 0.08 | 0.611 |
|  | cg16058600 | 0.10 | 0.515 | cg16246410 | 0.07 | 0.675 |
|  | cg11399967 | 0.08 | 0.590 | cg14617996 | 0.06 | 0.681 |
|  | cg25247969 | 0.07 | 0.656 | cg11358945 | 0.05 | 0.735 |
|  | cg07646377 | 0.06 | 0.697 | cg20804199 | 0.05 | 0.760 |
|  | cg11209121 | 0.06 | 0.714 | cg09906382 | 0.03 | 0.839 |
|  | cg16151636 | 0.06 | 0.724 | cg11209121 | 0.03 | 0.861 |
|  | cg14811105 | 0.04 | 0.810 | cg13500877 | 0.02 | 0.872 |
|  | cg18881247 | 0.02 | 0.894 | cg03339537 | 0.02 | 0.905 |
|  | cg11358945 | 0.02 | 0.906 | cg07646377 | 0.02 | 0.911 |
|  | cg15699971 | 0.01 | 0.924 | cg16151636 | 0.01 | 0.935 |
|  | cg03339537 | 0.01 | 0.940 | cg27074041 | 0.01 | 0.957 |
|  | cg01123282 | 0.00 | 0.982 | cg21226516 | 0.00 | 0.992 |
|  | cg20570611 | -0.01 | 0.940 | cg06101180 | -0.01 | 0.939 |
|  | cg00240195 | -0.04 | 0.813 | cg18941818 | -0.02 | 0.901 |
|  | cg16246410 | -0.04 | 0.805 | cg00194126 | -0.02 | 0.879 |
|  | cg20398418 | -0.05 | 0.754 | cg03953709 | -0.06 | 0.706 |
|  | cg18941818 | -0.05 | 0.732 | cg00240195 | -0.09 | 0.583 |
|  | cg14617996 | -0.06 | 0.709 | cg23474904 | -0.09 | 0.575 |
|  | cg27074041 | -0.06 | 0.707 | cg17654050 | -0.10 | 0.529 |
|  | cg17654050 | -0.07 | 0.675 | cg09408520 | -0.11 | 0.492 |
|  | cg06101180 | -0.07 | 0.653 | cg13945301 | -0.11 | 0.462 |
|  | cg20945253 | -0.10 | 0.538 | cg11399967 | -0.13 | 0.387 |
|  | cg25127975 | -0.10 | 0.528 | cg18881247 | -0.15 | 0.339 |
|  | cg07108936 | -0.13 | 0.391 | cg11932911 | -0.16 | 0.294 |
|  | cg21226516 | -0.14 | 0.378 | cg21506655 | -0.17 | 0.259 |
|  | cg11932911 | -0.14 | 0.370 | cg01123282 | -0.18 | 0.255 |
|  | cg09408520 | -0.22 | 0.165 | cg25127975 | -0.23 | 0.130 |

Notes: Blue highlights a positive correlation, and pink highlights a negative correlation. ^*^Nominally significant (*p* < 0.05).

Abbreviations: NR4A2; nuclear receptor subfamily 4A2, DNAm; DNA methylation, EOD; Epigenetics of Delirium.

Supplementary Table 5 : Correlation of age and blood DNAm at 18 CpGs in the *NPAS4* gene compared between delirium cases vs non-delirium controls in the EOD cohort

| Gene | Delirium Cases (N=43) | | | Non Delirium Controls (N=44) | | |
| --- | --- | --- | --- | --- | --- | --- |
|  | CpG | r | p-value | CpG | r | p-value |
| *NPAS4* | cg11635197 | 0.25 | 0.104 | cg10087138 | 0.48 | 0.001^*^ |
|  | cg10087138 | 0.24 | 0.126 | cg23484234 | 0.24 | 0.113 |
|  | cg19568845 | 0.22 | 0.164 | cg11635197 | 0.14 | 0.366 |
|  | cg15273822 | 0.15 | 0.330 | cg14699728 | 0.13 | 0.392 |
|  | cg02055483 | 0.15 | 0.346 | cg25233308 | 0.06 | 0.709 |
|  | cg23667955 | 0.13 | 0.401 | cg19568845 | 0.05 | 0.762 |
|  | cg09835239 | 0.08 | 0.614 | cg09835239 | 0.03 | 0.863 |
|  | cg25233308 | 0.07 | 0.675 | cg08715791 | -0.03 | 0.868 |
|  | cg13215078 | 0.04 | 0.808 | cg15273822 | -0.04 | 0.789 |
|  | cg22134325 | 0.01 | 0.960 | cg20420783 | -0.05 | 0.738 |
|  | cg14699728 | -0.02 | 0.894 | cg23667955 | -0.06 | 0.710 |
|  | cg20420783 | -0.04 | 0.787 | cg24435401 | -0.06 | 0.702 |
|  | cg24435401 | -0.05 | 0.749 | cg25565192 | -0.07 | 0.663 |
|  | cg08715791 | -0.08 | 0.613 | cg04768203 | -0.07 | 0.635 |
|  | cg25565192 | -0.08 | 0.611 | cg02055483 | -0.07 | 0.632 |
|  | cg23484234 | -0.12 | 0.435 | cg14326225 | -0.08 | 0.602 |
|  | cg04768203 | -0.14 | 0.388 | cg22134325 | -0.11 | 0.496 |
|  | cg14326225 | -0.14 | 0.371 | cg13215078 | -0.16 | 0.303 |

Notes: Blue highlights a positive correlation, and pink highlights a negative correlation. ^*^Nominally significant (*p* < 0.05).

Abbreviations: NPAS4; neuronal Per-Arnt-Sim domain protein 4, DNAm; DNA methylation, EOD; Epigenetics of Delirium.
